# Dance or disappear: Strategic sexual signalling in female Peninsular rock agama

**DOI:** 10.1101/2023.08.30.555377

**Authors:** Aravind Sridharan, Swapna Lawrence, Kavita Isvaran

## Abstract

Sexual signalling has largely been studied in the context of indirect competition among males through displays and mate choice by females. Historically, sexual signalling in females was expected to be of limited consequence. However, there is growing evidence of sexual signalling in females and its value in the context of sexual selection. Here, we test how females strategize the use of sexual signals in a socially polygynous lizard, *Psammophilus dorsalis*. We evaluated key hypotheses for how females should modulate their signalling, including male quality and mate availability. We simulate male quality using artificial male models to individually tagged, and intensely monitored wild female lizards, measuring their strategic investment in sexual signalling. We found that females invest more in signalling towards high quality males and increased their investment towards the later part of the only breeding season. Contrary to what is typically expected in a polygynous mating system, females not only invest in costly and elaborate sexual signals, they also modulate their usage to maximise their benefits and minimise their costs. Even in polygynous mating systems, dispersed distribution of individuals can result in females experiencing limitation in mate availability, resulting in costly sexual signalling.

## Introduction

Sexual selection, through direct competition and mate choice, favours the evolution of elaborate traits (Andersson, 1994). Some of the elaborate traits in animals are directed at the opposite sex as signals to attract mates (Jones and Ratterman, 2009) (Jones and Ratterman, 2009). Such sexual signalling can be expensive and the benefits from it may not be consistent. Consequently, individuals are expected to vary their investment in sexual signalling depending on the payoffs accruing to them (Higham et al., 2012; Simmons, 2015). Such variation in sexual signalling and its causes are commonly studied in males, whereas studies of females are few despite growing evidence of sexual signalling in females, reported from diverse taxa (Clutton-Brock, 2007; LeBas and Marshall, 2000; Stockley and Bro-Jørgensen, 2011). In this study, we examined strategic sexual signalling in females of a polygynous lizard.

Research on males suggests that variation in sexual signalling can arise as a result of males strategically modulating their signalling depending on the costs and benefits, they experience. Benefits are typically increased mating success and costs associated include energy expenditure and risk of predation. For example, lesser wax moth decrease their sexual signalling with an increase in the intensity of predation by bats (Edomwande and Barbosa, 2020). Energetic costs associated with sexual signalling can be high. Male wolf spiders *Hygrolycosa rubrofasciata* perform sexual signalling through drumming, which causes high metabolic rate and also results in increased mortality (Kotiaho et al., 1998).

What benefits might females gain from sexual signalling given that they too experience similar costs, including predation risk and energetic expenditure? It has been suggested that sexual signals can be best explained by females indicating their reproductive status in order to facilitate mating when they are receptive, while avoiding unnecessary coercion by males when they are not. If this is the case, we must expect relatively simple signals that can convey this information. However, reproductive status alone cannot explain the expensive investments females make on signals. For example, primates from the old world and apes such as chimpanzees display large red-coloured sexual swellings when ovulating. Such swellings are often very obvious and can weigh about 20-25 percent of their total body weight at their peak (Domb and Pagel, 2001).

Females can benefit by attracting the attention of high-quality males, in systems where mates are limited. For example, in monogamous mating systems with biparental care, the cost of desertion by males and females affects the reproductive success of their partner similarly (Jones and Hunter, 1993; Kraak and Bakker, 1998). Therefore, females are expected to signal to attract assessing males. Furthermore, in monogamous systems, females may compete through signalling to pair up with high quality fathers or for extra pair copulations with high quality males. Thus, the trade-offs associated with sexual signalling in females in monogamous systems is clear (Kokko and Johnstone, 2002). However, in polygynous systems, given that females have access to multiple males, what do they gain by investing in sexual signalling?

In polygynous mating systems, it is expected that females are the limiting sex and male fitness is maximised by mating with as many females as possible. Males should compete for females and mate indiscriminately. The maximum potential mating success is expected to be higher in males than in females (Orians, 1969). In contrast, theory predicts that females are not limited by mate availability but exercise mate choice for direct or indirect benefits. However, this assumption that females are not limited by access to mates may be strongly influenced by ecological conditions. In species that are spatially dispersed, males can be limited by economics of movement and are available to mate with females only within or close to the males’ home-range. Depending on how dispersed females are, the number of females within a male’s home range may be very low. Under such conditions, and given that male quality is likely to vary in a population, females may have limited access to high quality males at any given time. Consequently, females may benefit by using sexual signalling displays to attract high-quality males to mate with them or to settle in the area. Such displays can indicate the quality of the female (Edward and Chapman, 2011; Weiss, 2006). Males may then choose based on the quality or readiness to mate in females. This lack of access to high quality males driven by ecological conditions can explain the evolution of costly sexual signalling in females in polygynous systems, where historically sexual signalling on females was expected to be weak. Like in males, females should also vary their displays to maximise benefit and minimise cost. Our understanding of such strategic sexual signalling in female is poor, since very few studies have explored patterns and processes in sexual signalling in females of polygynous species (Amundsen et al., 1997; Higham et al., 2012; Huchard et al., 2009; LeBas and Marshall, 2000).

The Peninsular rock agama *Psammophilus dorsalis* is a very suitable model system to study the strategic use of signalling in females. In this socially polygynous, sexually dimorphic lizard species, both males and females use a range of displays, including dynamic colour change, postures and movement-based displays (Deodhar and Isvaran, 2018, 2017). Signals in the form of dynamic colour change and behavioural displays offer flexibility in when an individual invests in signalling. Peninsular rock agamas are dispersed within rocky outcrops of Peninsular India with males’ home-range typically overlapping with 2-3 females (Ranade and Isvaran, 2022). Typically, they have a lifespan of about one year and experience a single breeding season (Deodhar and Isvaran, 2017).

We tested multiple hypotheses that explore how females should vary their investment in sexual signalling. Since so little is known about sexual signalling in polygynous females, we focussed on key factors that sexual selection theory posits should affect fitness. We first considered male phenotype. Females are expected to accrue direct and/or indirect benefits by mating with high quality males (Andersson and Simmons, 2006; Clutton-Brock, 2007; Courtiol et al., 2016). Females should, hence, strategically vary their signalling in response to male quality, and invest more in their signalling effort towards high-quality males. Male agamas display their breeding season colours on rocks and boulders (henceforth perches) easily visible to females. Males vary in the colours they use (ranging from pale yellow to orange) when broadcasting signals to individuals in the vicinity (Deodhar and Isvaran, 2017). Given that orange colour males are most conspicuous to predators (Amdekar and Thaker, 2019), spending a greater amount of time in orange colour can indicate quality. We predict that females respond with more signalling towards orange-coloured males.

The second hypothesis we focussed on concerns mate availability. Access to multiple males, and especially high-quality males, in spatially dispersed systems can vary between females. *P. dorsalis* females typically encounter only one male at any given time within their home-range, and two males on average (with a range of 1-4) over their lifespan (Ranade and Isvaran, 2022). Females can strategize their investment in signalling to attract high-quality males. We hypothesise that females should vary their signalling depending on their access to high-quality males, with greater signalling by females experiencing lower mate availability. We predict that females found in areas of lower male densities signal more to a new male.

Finally, we examined terminal investment. Studies show that investment in sexual signalling can vary due to life history factors (Descamps et al., 2007; Mysterud et al., 2005; Velando et al., 2006). A key explanation in investment in costly traits is the balance between current and future reproductive fitness (Clutton-Brock, 1984). Since *P. dorsalis* typically experience one breeding season, we tested whether females signal more towards the latter part of the breeding season which represents the end of their reproductive lifespan.

We tested these hypotheses through field experiments with individually tagged wild lizards in natural ecological and social contexts. Variation in male quality was simulated using artificial male models. Local densities and the social environment of focal females was quantified by intense tagging and monitoring of lizards in the study area throughout the breeding season. This study explores how wild populations of a polygynous lizard species strategically vary their sexual signalling in different ecological and social contexts.

### Study system

*Psammophilus dorsalis* is a species of diurnal lizard with a socially polygynous mating system found on rocky outcrops throughout Peninsular India (Deodhar and Isvaran, 2017; Ranade and Isvaran, 2022). This species exhibits a strong male-biased sexual size dimorphism and sexual dichromatism. Over 90% of the population only live up to one year (Deodhar and Isvaran, 2017). Individuals communicate through visual signals such as movements, posture and colours (Deodhar and Isvaran, 2018; Ranade, 2020). Males broadcast in orange dorsally with black on the sides during active courtship interactions. However, males are commonly found to be in yellow dorsally and black on the sides through most of the breeding season. Male lizards also adopt a cryptic non-breeding colouration with pale yellow dorsally and grey body within the breeding season. Orange-black colouration observed during the breeding season is most conspicuous to predators (Amdekar, 2020). Female lizards display using postures such as gape, gular extension, movements such as head bob, and dynamic colour change in body colour from grey to dark body and red head (Ranade, 2020). Both males and females can change colour rapidly. Breeding season occurs from May to October (Deodhar and Isvaran, 2017).

#### Study site and data collection

The study was conducted in sheet rocks in and around Rishi Valley School in Chittoor district, Andhra Pradesh (13°32’ N, 78°28’ E). Typically, lizards were found in higher densities in rocky outcrops with boulders. Such areas were identified, mapped and data collection was carried out in them. Lizards were tagged and were regularly surveyed through their lifetime. Behavioural assays were carried out on such tagged lizards. This study was conducted between the years 2019 and 2022.

#### Tagging and surveys

Lizards were captured and tagged with a unique combination of ceramic beads for individual identification (more details in (Horta and Young, 2014). The lizards, once tagged with beads, are released back to the same site at which they were caught. The tagging operation was conducted complying with the guidelines of the institutional animal ethics committee (Indian Institute of Science).

Lizards were followed and observed through regular surveys. The surveys involve the observer walking along a fixed path through the sheet rocks and making note of all the lizards in the study area. Tagged individuals can be identified with images shot from a camera without disturbing the lizard. The lizards were noted down onto a map, which was later geo-referenced to get the exact location of the lizard. Colour status of males and gravidity status of females was noted for each lizard. Male colour states observed include orange-black, yellow-black, pale-yellow, pale (cryptic) and yellow-red colour states (See study system for more description). Each area was surveyed twice within the same session in order to maximise opportunistic sightings, however only tagged lizards were noted the second time. The survey data on lizard distributions were later used to analyse mate availability for focal females.

**Fig 1:**
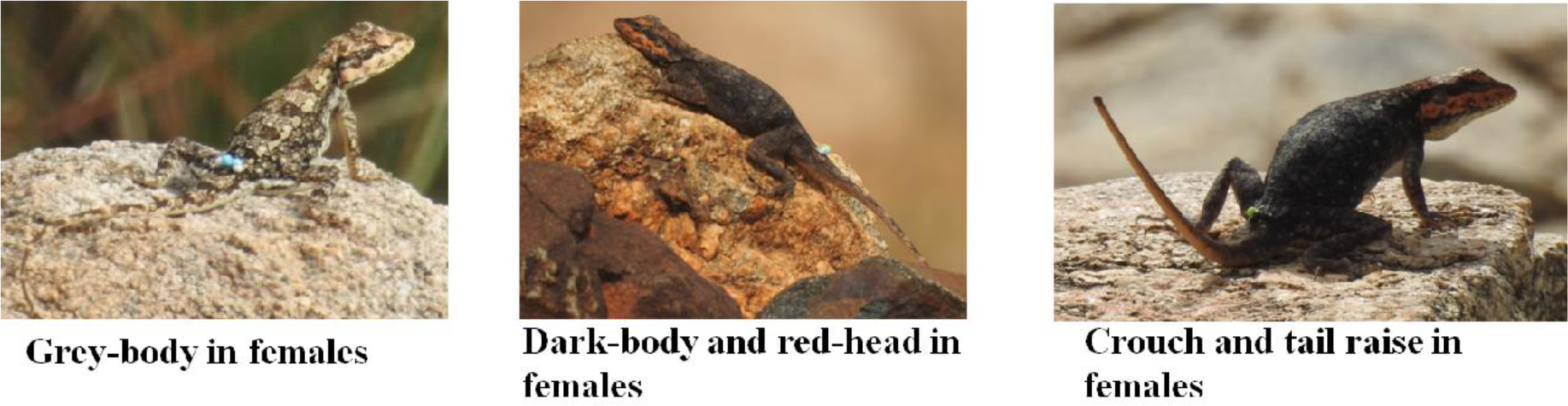
Different colour states and postural displays of female Psammophilus dorsalis

**Fig 2:**
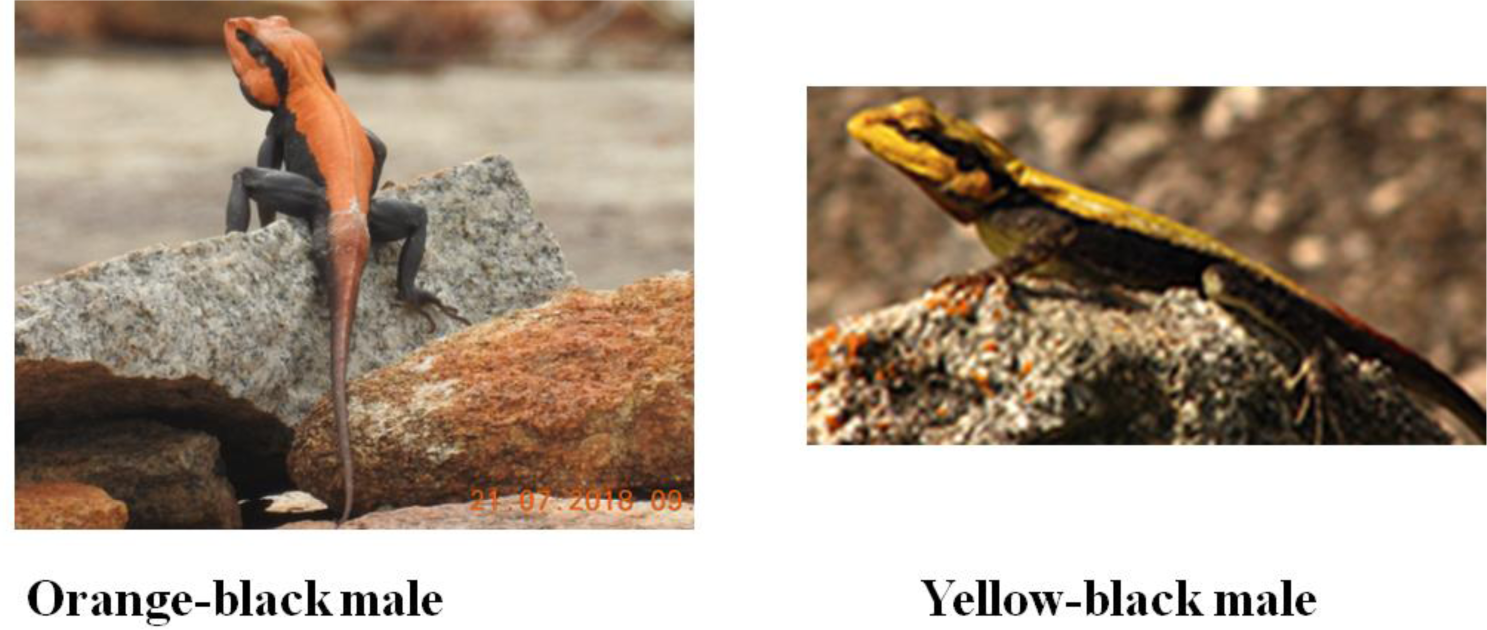
Colour states of male lizards in the breeding season

#### Model presentation assays

Female signalling responses to potential mates were assessed through field experiments on individually tagged females. 3D printed models mimicking male lizards (SVL-13 cm) were painted in three different colour states: yellow, and bright orange, and used in the model presentation assays. Female lizards were presented with these models at different times during their breeding lifespan. Both the colour states were commonly seen in the breeding season Model presentation observations were recorded on a video camera mounted on a tripod. The video camera is placed at a distance exceeding 15 metres from the focal lizard to avoid any disturbance to the focal lizard. Focal animal sampling (Altmann, 1974) was conducted on the tagged lizard as part of the initial observations for 15 minutes. This is followed by a presentation of the lizard model. The order in which the models (orange-black, yellow-black, pale-yellow) are presented to a tagged lizard was randomised. A gap of a minimum of 4 days was given between different model trials to avoid habituation to models by the focal lizard. The male model is carried towards the focal individual and placed on a perch at a distance less than 5 metres from the focal lizard. After placing the model typically on a perch where male lizards usually display, the observer returns back to record observations. We begin the experiment only when the model is in direct line of sight of the lizard. Observations of the focal lizard are carefully recorded on the video camera for a period of up to 30 minutes. Any adult conspecific lizard appearing within 5 m of the focal lizard during the experiment was noted. If the model or focal lizard was disturbed by other lizards in the area, then the trial was aborted. There were at least two individual models printed and painted of each type to offset the effect of the model. An elaborate ethogram of all the behaviours was created. We identified sexual signalling behaviours as those associated with courtship interactions from previous studies (Ranade, 2020) and ad lib observations of courtship behaviour in the field.

#### Data Analysis

The spatial information from regular surveys was geo-referenced from maps with the help of Qgis software. Model presentation assays were transcribed using the software BORIS (Friard and Gamba, 2016). We developed an ethogram with the help of previous studies and added relevant new behaviours. Frequency of short-duration behaviours and proportion of time spent in behavioural states were obtained and the rest of the analysis was carried out in R (Version 4.1.3)(Team, 2021).

#### Measuring social condition of lizards

The tagged female lizards we sampled through assays were followed through regular surveys. We considered survey data 30 days prior to the day of experiment to measure the provisional mate availability. Lizards were usually recorded multiple times in this period within this time. To each spatially marked sighting of a focal female, we used a 20 m buffer (using gbuffer function in R, package rgeos) and recorded the number of males that were observed for each unique sighting of the focal lizard. We measured the number of unique males each female was exposed to, in the 30-day period before the assay. For each sighting of the lizard, in the surveys, we were able to assign a total number of males and since the males were tagged, we were able to identify the number of unique males a female was exposed to, and this was considered as a proxy for provisional-mate availability.

#### Breeding season

The breeding season lasts from May till October. Experiments were predominantly carried out between June and October. June and July were considered as early breeding season and August-September were considered late breeding season.

#### Statistical Analyses

All analysis was carried out using R (Version 4.1.3). We calculated rates of behavioural events and proportion of time spent in behavioural states. To assess any co-variation in display behaviour we performed a principal component analysis on the data set. Since behaviours were not strongly correlated, indicating a possibility of different contexts and selection pressures, we considered each individual behaviour as an individual response variable. We then used generalised linear mixed effects models to test how female signalling varies in relation to male phenotype, mate availability and time during the breeding season (terminal investment). The response variables we considered were dark-body, red-head, tail raise and crouch posture. The predictor variables we examined were model colour (Colour states: Orange-black, yellow-black and pale), no of unique males found in the vicinity, and time in the breeding season (early/late). The number of males present in the vicinity was a discrete variable ranging from 0 to 8. Because there were relatively few samples for 2-8 (n < 10) we considered this as a categorical variable with three categories namely, 0,1 and greater than 1. Since crouch posture was observed in relatively few samples, we considered presence or absence of this behaviour in the observation sessions. The other three response variables (dark-body, red-head, tail raise) were represented as the proportion of time spent in that behavioural state. We fitted a zero inflated, beta-regression, mixed-effects model for each of these variables. Individual lizards were sampled multiple times according to their availability, therefore the inherent variation in behaviours of individual lizards was incorporated as a random effect in all the GLMMs described above. The package glmmTMB (Brooks et al., 2017) was used to fit GLMMs. Residuals were inspected to assess whether statistical model assumptions were satisfied.

## Result

We presented models 74 times (44 - orange and 30 - yellow) to 48 unique female lizards. Upon transcribing videos of our model presentation trials, we found that females show an array of behaviour; we shortlisted behaviour associated with male-female (courtship) interactions. Interestingly, most of these behaviours appear exclusive to a mating context since they do not overlap with behaviour described in the other common sexual selection context, namely female-female competition (Ranade et al., 2022). The final set of behaviours we examined were two colour states and two posture-based displays (Fig 3). We analysed female signalling response to male phenotype, mate availability and time during the breeding season within the common framework of GLMMs.

**Figure 3:**
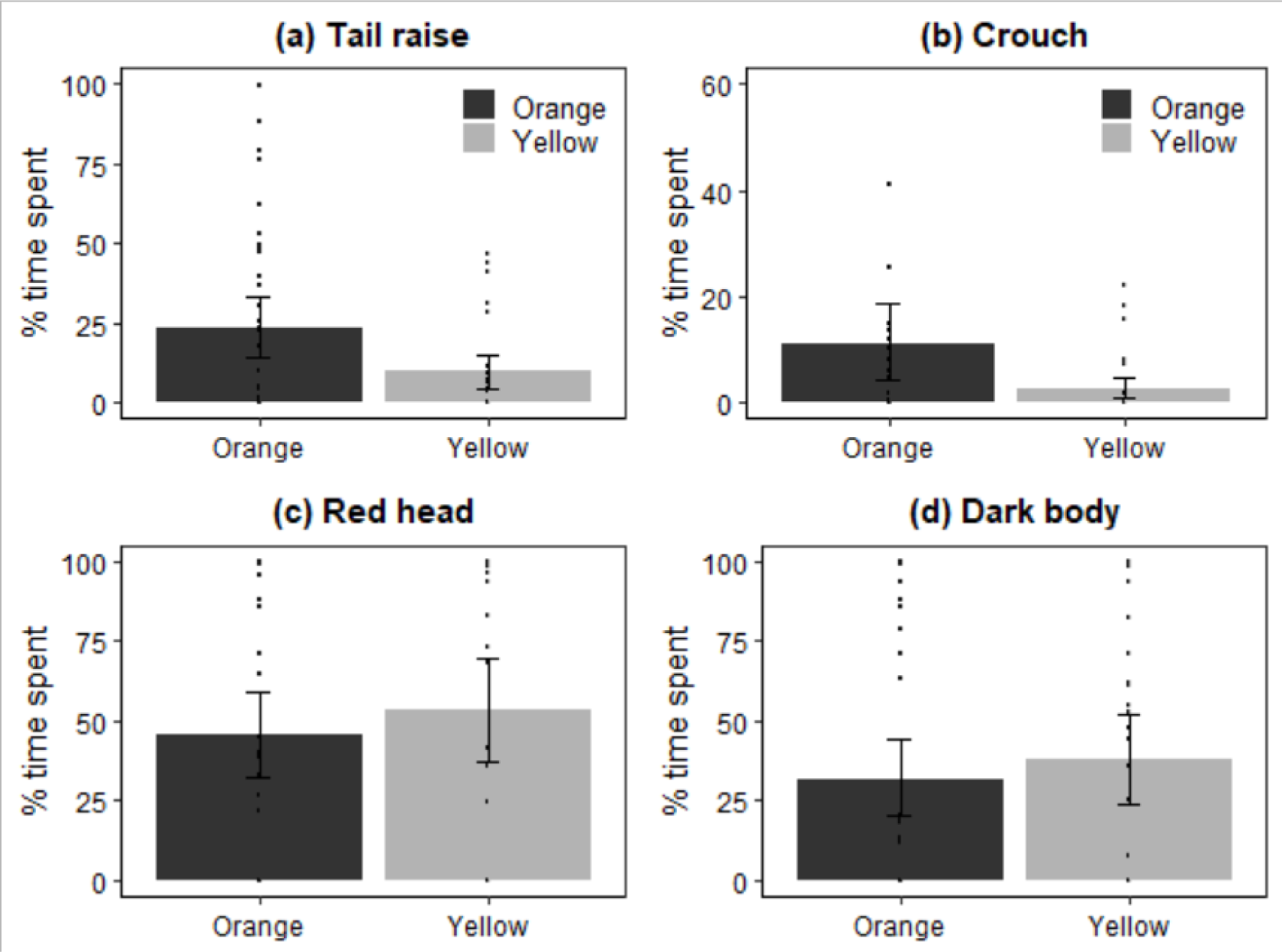
Comparison between percentage time spent in (a) tail raise, (b) crouch, (c) red head, (d) dark body, in response to models in orange (black bars) and yellow (grey bars) colour states. Error bars represent 95% bootstrapped confidence intervals around the mean.

The degree of signalling by females varied in response to male phenotype and to time during the mating season. Females varied their signalling through postures, tail raise and crouch, in response to the colour state of the male model (Fig 3, Table 1). Females spent more than twice the amount of time in tail raise, on average, when presented with an orange male model than when presented with a yellow model. The use of crouch posture increased by 5 times in the presence of orange male model compared with yellow male model. However, females did not vary their colour states detectably in response to male phenotype (Fig 3 Table 1).

**Table 1:** Results from models examining variation in proportion of time spent (Tail raise, crouch, red-head and dark body) on simulation of yellow model and orange model. Females appear to increase the proportion of time spent in tail raise in the presence of orange model. Females also show increase in tail raise, crouch and dark body through the later part of the breeding season. A trend of increased signalling of crouch was observed in the presence of orange model. Zero-inflated GLMMs with beta regression family were used for tail raise, red head and dark body. GLMMs with binomial distribution was used for crouch posture. Estimates, 95% confidence intervals (CI) and results from likelihood ratio tests are shown. Additionally, model coefficients were bootstrapped with 5000 iterations and confidence intervals are shown.

| Tail Raise |  |  |  |  |  |  |  |  |
| --- | --- | --- | --- | --- | --- | --- | --- | --- |
| | Estimate | Confidence Interval | | $\chi^2$ | df | p | Bootstrapped CI | |
|  |  | 2.5% | 97.5% |  |  |  | 2.5% | 97.5% |
| Intercept | -0.185 | -0.979 | 0.608 |  |  | 0.646 | -1.312 | 0.869 |
| **Yellow model** | **-0.858** | **-1.755** | **0.039** | **3.203** | **1** | **0.06** | -1.764 | 0.328 |
| No of male 1 | -0.490 | -1.554 | 0.573 | 1.016 | 2 | 0.365 | -2.025 | 0.510 |
| No of male >1 | -0.346 | -1.346 | 0.654 | 1.016 | 2 | 0.497 | -1.762 | 1.069 |
| **Late breeding season** | **0.863** | **-0.002** | **1.729** | **4.070** | **1** | **0.05** | -0.054 | 2.038 |
| ZI. Intercept | **0.05407** | **-0.402** | **0.51** |  |  | 0.816 |  |  |

| Crouch Posture |  |  |  |  |  |  |  |  |
| --- | --- | --- | --- | --- | --- | --- | --- | --- |
| | Estimate | Confidence Interval | | $\chi^2$ | df | p | Bootstrapped CI | |
|  |  | 2.5% | 97.5% |  |  |  | 2.50% | 97.5% |
| Intercept | **0.215** | **0.015** | **0.417** |  |  | **0.035** | 0.007 | 0.428 |
| Yellow model | -0.162 | -0.348 | 0.023 | 0.097 | 1 | 0.085 | -0.382 | 0.086 |
| No of male 1 | 0.153 | -0.098 | 0.405 | 0.490 | 2 | 0.231 | -0.131 | 0.562 |
| No of male >1 | 0.051 | -0.191 | 0.295 | 0.490 | 2 | 0.675 | -0.313 | 0.210 |
| **Late breeding season** | **0.268** | **0.050** | **0.488** | **0.02** | **1** | **0.016** | -0.024 | 0.534 |

| Red Head |  |  |  |  |  |  |  |  |
| --- | --- | --- | --- | --- | --- | --- | --- | --- |
| | Estimate | Confidence Interval | | $\chi^2$ | df | p | Bootstrapped CI | |
|  |  | 2.5% | 97.5% |  |  |  | 2.5% | 97.5% |
| Intercept | 1.250 | 0.467 | 2.034 |  |  | 0.001 | 0.515 | 2.028 |
| Yellow model | 0.097 | -0.625 | 0.82 | 0.79 | 1 | 0.79 | -0.704 | 1.231 |
| No of male 1 | 0.066 | -0.766 | 0.9 | 0.593 | 2 | 0.875 | -0.987 | 0.815 |
| No of male >1 | 0.411 | -0.397 | 1.219 | 0.593 | 2 | 0.318 | -0.328 | 1.198 |
| Late breeding season | 0.217 | -0.465 | 0.9 | 0.532 | 1 | 0.532 | -0.278 | 0.943 |
| ZI. Intercept | -0.383 | -0.847 | 0.081 |  |  | 0.106 |  |  |

| Dark Body |  |  |  |  |  |  |  |  |
| --- | --- | --- | --- | --- | --- | --- | --- | --- |
| | Estimate | Confidence Interval | | $\chi^2$ | df | p | Bootstrapped CI | |
|  |  | 2.5% | 97.5% |  |  |  | 2.5% | 97.5% |
| Intercept | 0.871 | 0.071 | 1.673 |  |  | 0.032 | -0.142 | 1.664 |
| Yellow model | -0.300 | -1.097 | 0.497 | 0.461 | 1 | 0.46 | -1.169 | 0.973 |
| No of male 1 | -0.328 | -1.288 | 0.631 | 0.276 | 2 | 0.501 | -1.543 | 0.641 |
| No of male >1 | -0.736 | -1.627 | 0.154 | 0.276 | 2 | 0.105 | -1.680 | 0.293 |
| **Late breeding season** | **0.828** | **0.057** | **1.601** | **0.038** | **1** | **0.035** | 0.169 | 1.808 |
| ZI. Intercept | -0.054 | -0.510 | 0.402 |  |  | 0.816 |  |  |

Sexual signalling in females did not show a statistical trend with respect to mate availability (Fig 4) but showed clear differences across the breeding season. Females increased sexual signalling in the later part of the breeding season ie: August and September. (Fig 5; Table 1) compared with the early breeding season. Dark body, tail raise and crouch increased in the later part of the breeding season. (Table 1). Use of sexual signals such as dark-body, tail raise and crouch at least doubled in their response to orange coloured male model (Fig 3).

**Figure 4:**
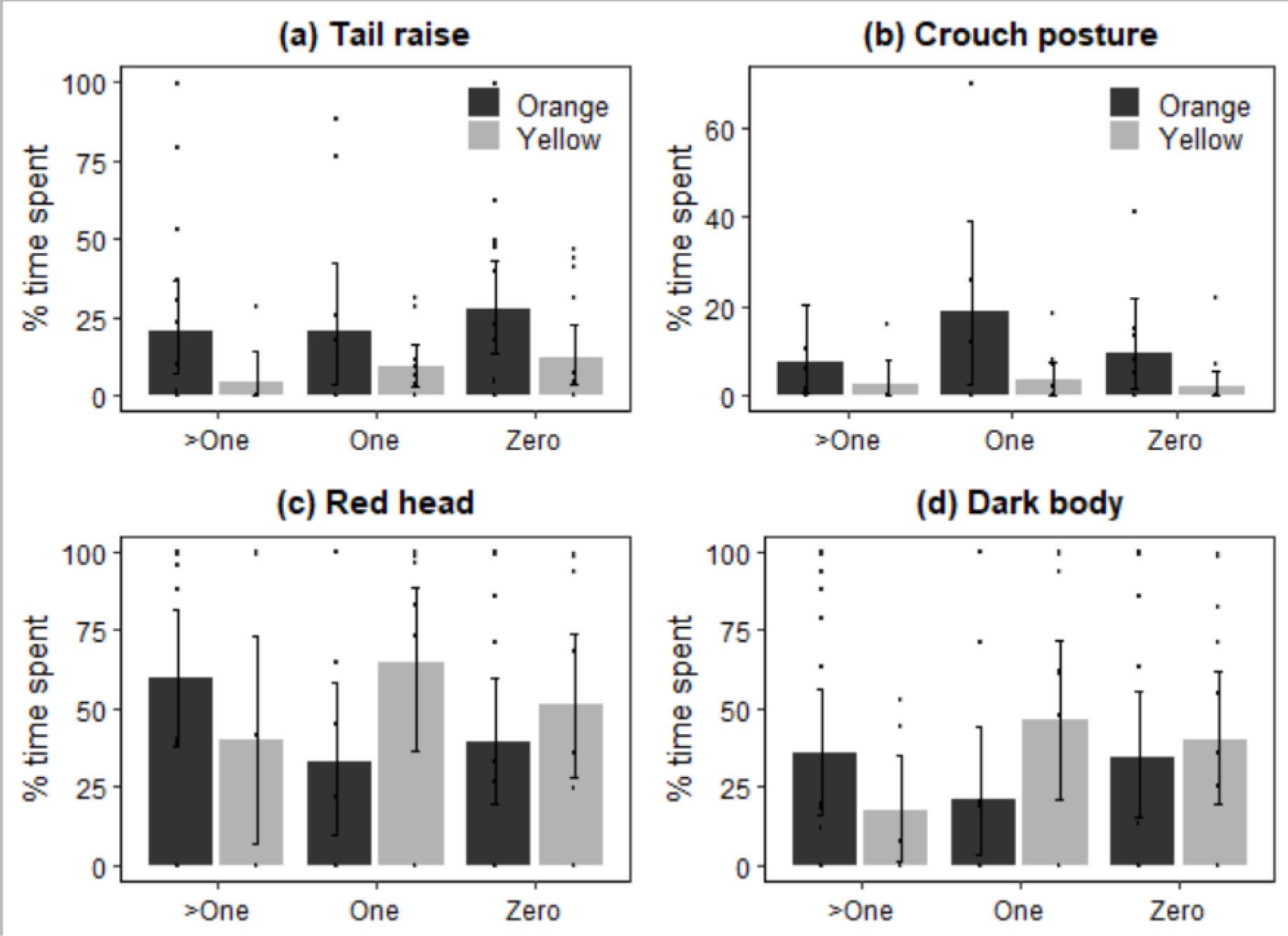
Comparison between percentage time spent in (a) tail raise, (b) crouch, (c) red head, (d) dark body by female lizards, in response to orange (black bars) and yellow (grey bars) models with varying access to male lizards (Zero, one and greater than one). Error bars represent 95 percent bootstrapped confidence intervals around the mean.

**Figure 5:**
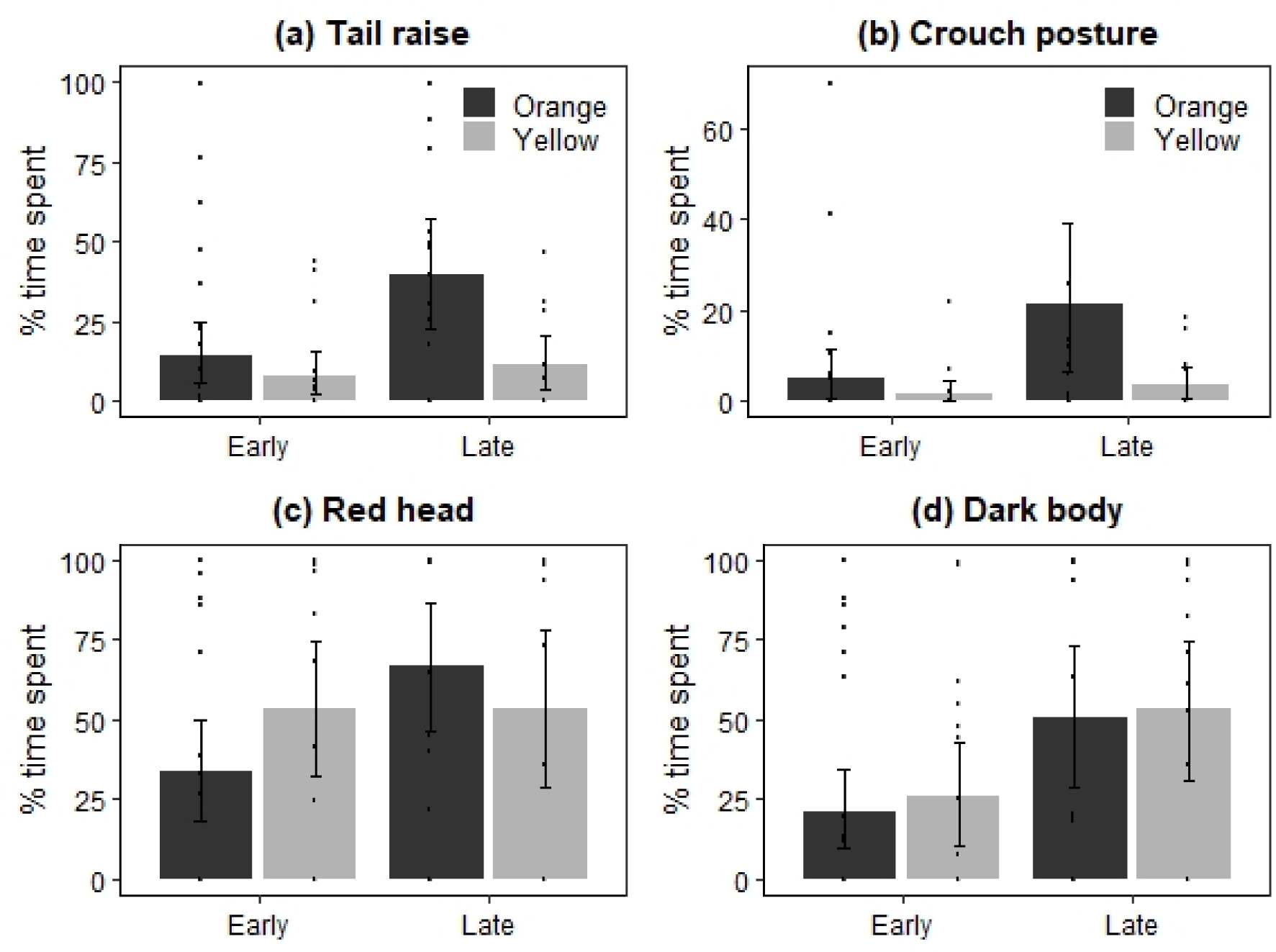
Comparison between percentage time spent in (a) tail raise, (b) crouch, (c) red head, (d) dark body by female lizards, in response to orange (black bars) and yellow (grey bars) models in early and late breeding season. Error bars represent 95 percent bootstrapped confidence intervals around the mean.

## Discussion

Through field experiments we show that female Peninsular rock agama strategically vary their signalling to attract mates. This detailed study is the first of its kind to examine how females in a polygynous mating system strategically modulate their sexual signals. Females use diverse posture-based and dynamic colour-change based signals to signal to potential mates. As predicted by the male phenotype hypothesis, females invested more in sexual signals towards experimentally simulated high-quality males. We argue that contradicting popular expectations from sexual selection theory, the usage of costly displays to attract high-quality males clearly indicates that access to such males is likely to be limited in this socially polygynous system.

Why should females attempt to attract high quality males? We propose that the answer may lie in how ecology of male and female dispersion may affect operational sex ratio (Emlen and Oring, 1977). In *P. dorsalis* males defend resources such as perch and nesting areas which are essential to females. Territoriality in females, perhaps as a result of uneven distribution of resources, has them dispersed in the rocky outcrops. Males defend multiple territories of females, and are dispersed in larger home ranges. When territorial males defend females, the cost of invading other male territories and economics of movement can limit their access largely to the females within their home range. Furthermore, males may only be able to economically defend a few females, depending on how dispersed they are. In *P. dorsalis*, males typically overlap the home ranges of only 2 females (interquartile range = 1–4; Ranade & Isvaran 2022). Consequently, the realised operational sex ratio can be lower than what is usually predicted resulting in females experiencing limitation in access to high quality males.

Previous research has shown that male Peninsular rock agama exhibit individual variation in the time spent on mate-attracting signals, such as orange-black coloration (Deodhar and Isvaran, 2017). On an average, females share their home range with less than two males during the breeding season at a given point in time (Ranade and Isvaran, 2022). This potentially indicates variability in access to high quality males. Therefore, females are likely to benefit immensely by indirectly competing with each other through sexual signalling in order to attract males. The dispersion pattern shown by P. dorsalis is commonly seen in other animals (Brown and Orians, 1970), suggesting that such female signalling in response to limited mate availability in socially polygynous species may be widespread.

In dispersed species with territorial males and females, the availability of high-quality males can vary for each female. To gain a better understanding of mate availability for females, we examined the social conditions of female lizards. Although we expected females to increase their sexual signalling when limited by access to high quality males, we found little difference in how females signal to males in different mate availability contexts. Even though we hypothesised that females are expected to decrease their investment in sexual signalling with increased mate availability, our understanding of the rate at which such investment reduces remains unclear. We had to categorise available males to zero, one and greater than one due to sample size constraints and large range variation in mate availability towards different females. Our categorization may not be refined enough to illustrate the difference in their response.

Our results also indicate that females adjust their investment in sexual signalling over their adult life. According to the terminal mating hypothesis, as the prospect of survival decreases for an organism, they are likely to invest more towards reproductive effort (Clutton-Brock, 1984). Although females lay multiple clutches of eggs within one breeding season, *P. dorsalis* live up to only breeding season (Deodhar and Isvaran, 2017). As a result, females experience the final opportunity to attract mates in the later part of the breeding season. “The increase in signalling by females towards the late breeding season indicates that females balance current and future investment over their lifespan. Females optimising their reproductive success through such strategies in fecundity and parental care traits has been previously reported in an array of taxa (Mysterud et al., 2005; Velando et al., 2006), especially with multiple breeding tenures. Our work indicates that such optimisation may also act on mating traits in females.

Studies on sexual selection often focus on competition in males and choosiness in females (Andersson and Simmons, 2006). *P. dorsalis* is indeed an excellent model system to study such male competition and female choice because of the conspicuous colour in males and male-biased sexual size dimorphism (Ranade and Isvaran, 2022). However, our study presents clear support for indirect competition in females through strategic sexual signalling. There is increasing evidence of male mate choice even in species which are not role reversed (Amundsen and Forsgren, 2001; Edward and Chapman, 2011). Our findings support the need to consider perspectives on mutual mate choice in such systems.

While theoretical understanding of male mate choice has received some attention (Edward and Chapman, 2011), empirical studies that show how males benefit from assessing females are relatively few. The benefits males acquire have been reported in primates, where size of sexual swellings in females has been an indicator of offspring quality (Domb and Pagel, 2001; Huchard et al., 2009). Our study presents an instance of a female using costly sexual signals and strategically varying them; these findings point to the need for the study of male strategies in assessing such signals and the benefits gained through male mate choice.

It is worth noting that sexual signalling in females, even when overt, can often be hidden unless the right stimuli is present and may be overlooked by researchers. However, such sexual signalling can significantly influence the behaviour and ecology of a species. Experimental manipulation in wild conditions clearly proves to be very important in evoking sexual signalling responses, and our study represents one of the first instances where such responses were empirically tested.

In conclusion, our field experiments in a wild lizard population, which involved simulating different mate quality by presenting male models in different colour states, demonstrated the complex strategy employed by female Peninsular rock agama in the context of indirect competition. Females use a wide range of visual signals according to mate quality and they balance their usage through the breeding season. These findings challenge the typical expectations of a polygynous mating system, highlighting that females not only invest in costly and elaborate sexual signals, but they also modulate their usage to maximise their benefits and minimise their costs. We propose that even in polygynous mating systems, male and female dispersion patterns, influenced by ecological conditions, can result in females experiencing limitation in mate availability, resulting in costly sexual signalling.

## Acknowledgement

We wish to thank IISc for providing institutional facilities and Rishi Valley school and Nature Conservation Foundation for logistic support during data collection. We are grateful towards P. Somnath and interns (Aneesh Konar, Keren Pereira, Vipin Sonu, Vandana Kannan, Swathi Ramesh, Jabili Choudhary, Tryambak Dasgupta, Abhinetri M, Ruthvik S P) for help with data collection. We would also like to thank Department of Biotechnology – Indian Institute of Science (DBT-IISc) partnership program and Department of Science and Technology Fund for Improvement of Science and Technology (DST-FIST) and Department of Science and Technology – Science and Engineering Research Board (DST-SERB) grant - CRG/2018/003928 for funding this research, and the Ministry of Human Resource Development (MHRD) for providing fellowship during the course of study. The capturing and handling of animals used in the study was approved by the Institutional Animal Ethics Committee constituted by the Indian Institute of Science (CAF/ Ethics/605/2018).

